# An Adaptive Cue Selection Model of Allocentric Spatial Reorientation

**DOI:** 10.1101/860031

**Authors:** James Negen, Laura Bird, Marko Nardini

**Affiliations:** Durham University, Department of Psychology

## Abstract

After becoming disoriented, an organism must use the local environment to reorient and recover vectors to important locations. Debates over how this happens have been extensive. A new theory, Adaptive Combination, suggests that the information from different spatial cues are combined with Bayesian efficiency. To test this further, we modified the standard reorientation paradigm to be more amenable to Bayesian cue combination analyses while still requiring reorientation, still requiring participants to recall goal locations from memory, and focusing on situations that require the use of the allocentric (world-based; not egocentric) frame. 12 adults and 20 children at 5-7 years old were asked to recall locations in a virtual environment after a disorientation. They could use either a pair of landmarks at the North and South, a pair at the East and West, or both. Results were not consistent with Adaptive Combination. Instead, they are consistent with the use of the most useful (nearest) single landmark in isolation. We term this Adaptive Selection. Experiment 2 suggests that adults also use the Adaptive Selection method when they are not disoriented but still required to use a local allocentric frame. This suggests that the process of recalling a location in the allocentric frame is typically guided by the single most useful landmark, rather than a Bayesian combination of landmarks – regardless of whether the use of the allocentric frame is forced by disorientation or another method. These failures to benefit from a Bayesian strategy accord with the broad idea that there are important limits to Bayesian theories of the cognition, particularly for complex tasks such as allocentric recall.

---

Reorientation is the process of recovering one’s heading and position in a given space. This is a process that allows a disoriented organism to recover the correct vector to important locations. The ability to do this is a key adaptation for the vast majority of mobile organisms. The study of how humans and other animals do this has moved forward our understanding of both cognition (Lee, 2017; Mou & Zhou, 2013; Nardini, Thomas, Knowland, Braddick, & Atkinson, 2009; Twyman, Holden, & Newcombe, 2018) and the mammalian brain (Cressant, Muller, & Poucet, 1997; Ito, Zhang, Witter, Moser, & Moser, 2015; Keinath, Julian, Epstein, & Muzzio, 2017; Park, Brady, Greene, & Oliva, 2011). This has especially become a crucial point in developmental studies of spatial cognition, igniting a debate over modular cognition (Cheng, 1986; Doeller & Burgess, 2008; Hermer & Spelke, 1996, 1994) versus adaptive behaviour (Cheng, Huttenlocher, & Newcombe, 2013; Learmonth, Nadel, & Newcombe, 2002; Ratliff & Newcombe, 2008; Twyman et al., 2018). A recent paper formalizes and details a specific proposal concerning adaptive behaviour (Xu, Regier, & Newcombe, 2017). More than just adaptive, this new theory posits that children’s use of different cues to reorient is fully rational and Bayesian. The full name of the model is the adaptive cue combination model of human spatial reorientation. For brevity, we will refer to it as Adaptive Combination. The present study seeks to further test this model.

Adaptive Combination is an important model for the study of developing spatial cognition. Despite decades of research (Cheng et al., 2013; Lee, 2017; Miller, 2009), there are still debates about the way that different cues are used by young children to reorient. For example, an early theory posited that reorientation only depends on environmental surfaces or boundaries, with the exception of adults who have a linguistic mechanism of incorporating additional information (Hermer & Spelke, 1994). In other words, if the target was to the right of a wall that was relatively short and coloured blue, an adult can synthesize the two pieces of information (right of short + right of blue) into one linguistic construct that could guide behaviour: ‘to the right of the short blue wall’. This theory, like many after it, faced a serious difficulty. It was discovered that young children’s performance can be improved through the addition of a non-boundary cue as long as the room is sufficiently large (Learmonth et al., 2002). This showed that the process is not purely dependent on boundary information, even in young children. The present paper seeks to test Adaptive Combination independently in the hopes of leading towards a consensus on how developing spatial cognition handles multiple reorientation cues.

If Adaptive Combination is true, it is also a breakthrough finding for the study of developing Bayes-like reasoning in perception and memory. Almost all previous studies to look at Bayesian cue combination in children under 10 years old have returned negative results (Adams, 2016; Burr & Gori, 2012; Chambers, Sokhey, Gaebler-Spira, & Kording, 2018; Dekker et al., 2015; Gori, Sandini, & Burr, 2012; Jovanovic & Drewing, 2014; Nardini, Bedford, & Mareschal, 2010; Nardini, Begus, & Mareschal, 2013; Petrini, Remark, Smith, & Nardini, 2014), including one that looked at combination of cues for spatial recall (Nardini, Jones, Bedford, & Braddick, 2008). For example, when judging a horizontal location with a spatialized audio cue and a brief visual cue, children under 10 years old fail to integrate the two efficiently; the precision of their judgements is not any better than with the visual cue alone (Gori et al., 2012). If the process of reorientation really does happen with full Bayesian efficiency, this means that spatial cognition is an exception to the general rule. Children might begin reasoning in a Bayes-like way in terms of reorienting first, then extend this to other cognitive processes throughout childhood. This again makes it vital to see if this theory can be verified independently: it has serious potential impact in terms of both spatial cognition and a general theory of how Bayesian reasoning develops.

The remaining sections of the Introduction (1) detail this model and give further terminology; (2) specify the gaps in evidence for Adaptive Combination; (3) explain key choices in the present study’s design; and (4) detail specific hypotheses and the way they will be tested.

## The Adaptive Combination Model and Terminology

First, we need to make it clear what the Adaptive Combination model is and how it works. Adaptive Combination is a Bayesian model of cue combination. In general terms, suppose that a participant is given one cue in isolation to do a given task. Let that process be governed by a function *f*_1_(Response|Target) that specifies how likely given responses are to given targets in the task. Suppose the same participant is given a different cue in isolation, now governed by *f*_2_(Response|Target). Suppose further that the function *f*_1+2_(Response|Target) governs behaviour when both cues are presented at the same time. By definition (Ernst & Banks, 2002), the process is a Bayesian cue combination process if and only if

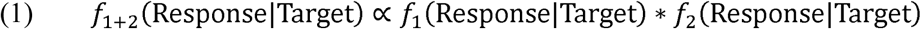

This is the fully rational process – the one that maximises the precision (minimising the variance) of responses with both cues available (Ernst & Banks, 2002).

An example might be helpful. Some more terminology will be needed. In the typical reorientation paradigm (Figure 1), children are placed in a rectangular arena and shown a target hidden in one corner. They are disoriented and then released to search one of the corners for the target. The correct corner is conventionally called C (for correct), the rotational equivalent called R (for rotational equivalent), the corner on the same short wall as the target called N (for near), and the corner on the same long wall called F (for far). If the geometry of the room is the only available cue, this is a G condition (for geometry). If there is also something unique about one of the walls to associate with the target, then it is an A+G condition (Associative + Geometry).

**Figure 1.**
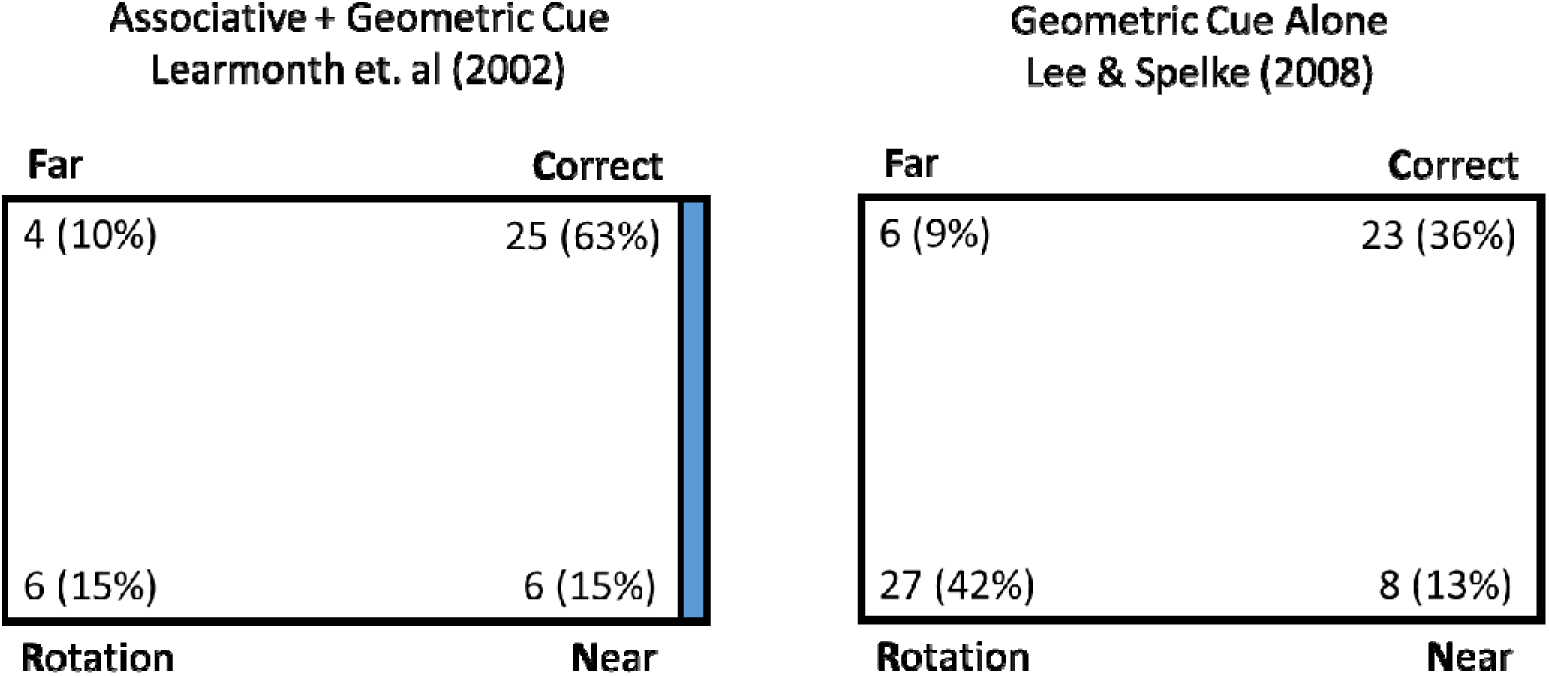
Examples of previous results with reorientation tasks. Children were placed in a rectangle arena with four hiding locations, one in each corner. The target was hidden in the corner marked ‘correct’. Children were first disoriented and then allowed to search for the target. On the right, participants can only use the geometry to find the target. This means that they respond in roughly equal numbers at the correct corner and its rotational equivalent (i.e. both corners with a long wall to the left and a short wall to the right). On the left, one of the walls was coloured blue, while the others were white. This associative cue made it possible to disambiguate the correct corner and its rotational equivalent. Children responded more often at the correct corner.

**Figure 1.**
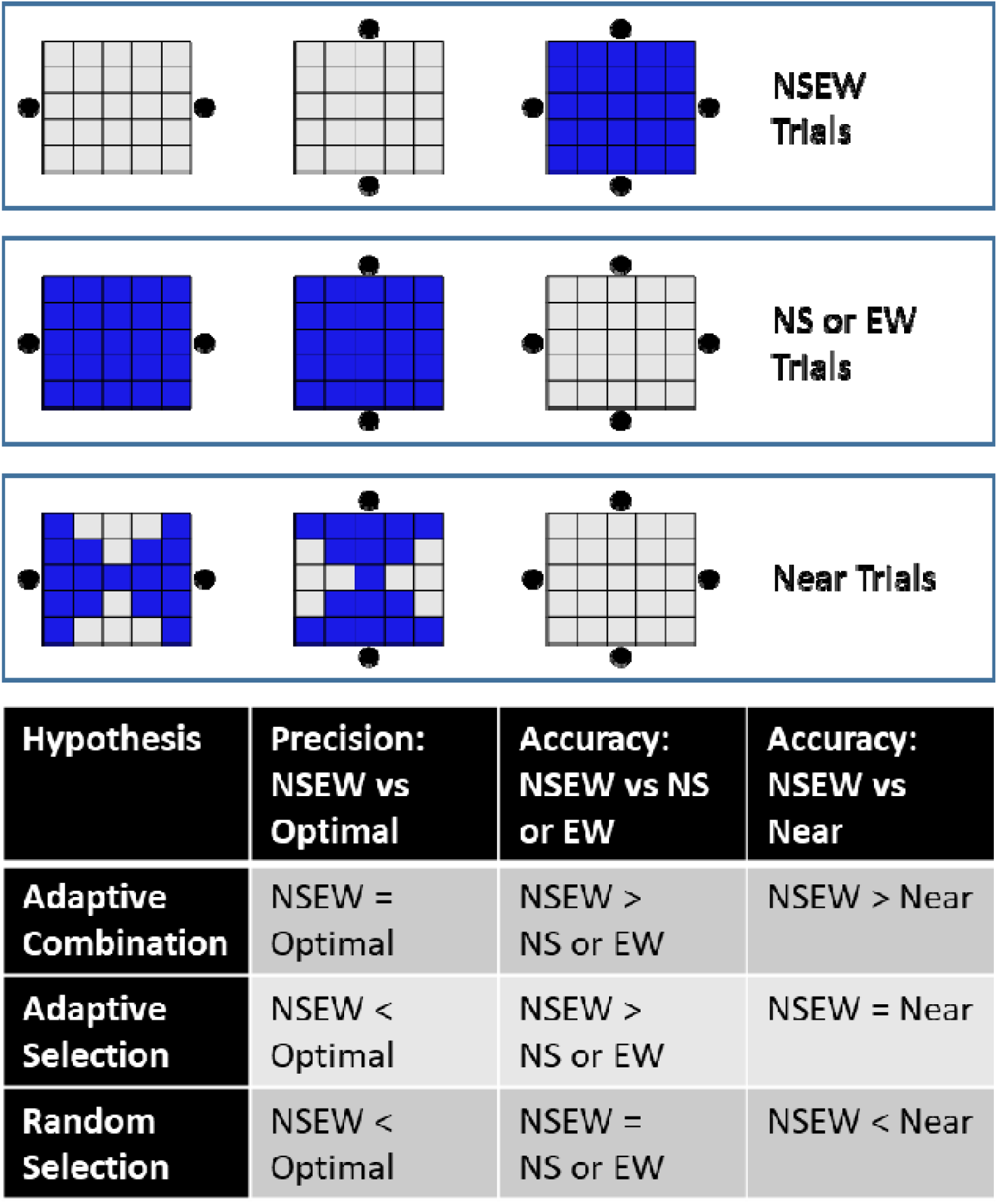
Trial types and key predictions. In terms of the methods, every trial is a NS trial (with North and South landmarks visible), an EW trial (East and West visible), or a NSEW trial (all four visible). However, for the analysis to differentiate between hypotheses, it is useful to regroup the trials. On the top half of this figure, the blue squares indicate which trials are included in each of the three regrouped categories. There are 25 possible targets in a 5×5 grid. The black dots indicate which landmarks are visible during those trials. NSEW includes all trials where all four landmarks were visible. NS or EW includes all trials where only the North and South landmarks were visible, plus the trials where only the East and West landmarks were visible. Near trials are a subset of NS or EW trials where the participant has the nearest possible landmark (or at least one of them if several are equidistant). The bottom table gives predictions. As the table shows, these regroupings allow us to test different predictions from the different hypotheses. The text justifies each entry in the lower table.

We can now insert some specific numbers and give example calculations. With boundary geometry alone, suppose participants respond at C 40% of the time, R 40%, N 10%, and F 10%. That is. Suppose that an associative cue alone would point a child to C 40%, R 10%, N 40%, and F 10%. That is. Assuming that Adaptive Combination is correct, we can now predict how often they will respond at each location during an A+G condition. We multiply to obtain P(C) = 0.4*0.4 = 0.16, P(R) = 0.04, P(N) = 0.04, and P(F) = 0.01. These then have to be normalized (divided by their sum) to arrive at the final probabilities. Those are P(C) = 64%, P(R) = 16%, P(N) = 16%, and P(F) = 4%. That is.

Equation 1 can lead to a variety of different interesting patterns, but one will be particularly critical here. In the example, the two cues presented together led to a higher proportion of correct answers (64%) than either cue alone (40%). In general, if both f_1_ and f_2_ have some kind of concentration (a discrete mode or a continuous peak) in the same place, then f_1+2_ will have an even greater concentration around the same place.

## The Need for Additional Scrutiny

Second, we need to clarify where the gaps in evidence for Adaptive Combination exist. In the literature on Bayesian perception and decision-making, there is a standard set of three findings that are used to show that two cues are combined in a Bayesian manner. This is routine enough that it has been codified into a tutorial with supporting R packages (Ernst, Rohde, & van Dam, 2016). The procedure measures how precise participants are with one cue in isolation, the other cue in isolation, and both cues together. It then must be shown that (1) precision is better with both cues versus the first cue in isolation; (2) precision is better with both cues versus the second cue in isolation; and (3) precision is not significantly different with both cues versus the Bayesian optimal prediction (predicated on Equation 1). These findings rule out the alternative hypothesis that either single cue is being used in isolation; otherwise, we would not expect better precision when both are presented. These findings also speak against the alternative hypothesis that the two cues are being used together in some non-Bayesian fashion; since Bayesian cue combination is the optimal way to improve precision, no other process could also match the optimal Bayesian precision.

Unfortunately, the paper arguing for Adaptive Combination (Xu et al., 2017) only provides one of the three pieces of evidence. Specifically, it reviews evidence that performance with A+G conditions is better than G conditions. It does not show that performance with A+G conditions is better than A conditions (where only an associative cue is presented; in practice, a square room with a single uniquely coloured wall). It also does not use data from G conditions and A conditions to derive predictions for A+G conditions and compare that to A+G data. This leaves open the alternative hypothesis that children may complete an A+G condition by only using the associative cue.

We tried to fill this gap as best as possible by looking through the available literature. Unfortunately, this attempt failed to show that performance in A+G conditions is different than performance in A conditions. We re-examined previous data for an A+G condition (Learmonth et al., 2002) and an A condition (Hermer-Vazquez, Moffet, & Munkholm, 2001). Since results are known to depend on age, we used the data from 5 year olds from both studies. As Adaptive Combination is theorized to ignore associative cues in small rooms, rather than combine them, we also used the data from the larger room in the Learmonth et al. (2002) study. These data are reproduced in Table 1. Analysis suggests that the two distributions are not reliably different, *X*^2^(3) = 2.08, p = 0.56. Further, a Bayesian version of this analysis can test the hypothesis that the response distributions are the same versus the hypothesis that they are different. This analysis results in BF_01_ = 18.44, considered ‘strong’ evidence that they are actually the same. We would present additional analyses, but A conditions are relatively rare in the literature and this was the only comparison we could find with a sufficient age match, the standard methods described above, and a full report of the response distributions.

**Table 1.**

|  | Correct | Rotation | Near | Far |
| --- | --- | --- | --- | --- |
| H-V 2001 (A) | 65 (51%) | 21 (16%) | 27 (21%) | 15 (12%) |
| L 2002 (A+G) | 25 (63%) | 6 (15%) | 5 (13%) | 4 (10%) |

From the point of view of the literature on Bayesian perception and decision-making, this makes it clear that further evidence is needed for the Adaptive Combination model. The reanalysis of the available previous data suggests that Bayesian reasoning is not occurring here. Instead, it suggests that participants in an A+G condition are merely using the associative cue to complete the task. This is certainly an unusual interpretation – to our knowledge, it has not previously been tested if performance in an A+G condition might depend entirely on the use of the associative cue. However, it may still be possible to improve on this analysis. This will be described in more detail in the next section, but briefly: the number of trials per participant is (radically) smaller than most cue combination studies, it is not ideal to use cues that are not equally useful, and it is not ideal to use between-subjects data. We therefore designed a new study to test Adaptive Combination in a more rigorous fashion.

## The Present Study

Third, we now outline key design decisions for the study. To do this, we need to comment on our focus with this design. Many researchers are specifically interested in the way that reorienting organisms (human or non-human, child or adult) use geometric boundaries (Lee, 2017). That is not our focus here. Our motivation here is the way that Adaptive Combination seeks to make a single coherent and compact explanation of the method that organisms use to reorient – how they regain a sense of position and heading with respect to important locations around them. The Adaptive Combination model is described in a way that is very general to different cues; indeed, the authors apply it to explaining how non-rectangular enclosures, various associative cues, polarizing cues, and linguistic information given to the participant are combined. Landmarks are also mentioned in the narrative account of the study’s remit (Xu et al., 2017). Under our reading, it is designed to accommodate any cue that could be used to reorient under a single mathematical framework with a compact mathematical law (Equation 1). Further, it posits that this happens as a Bayesian cue combination process. Our goal in the present study was not to investigate specifically how people use rectangular enclosures, but instead to find a situation in which we could clearly distinguish whether or not multiple cues are used to recall goal locations in a Bayesian manner after disorientation. This is a test of the general form of Adaptive Combination, rather than a test of how it specifically applies to particular situations that appear frequently in the literature. In short, we are interested here in reorientation – not the typical reorientation paradigm.

Any cue combination study faces a number of routine problems to overcome (Ernst et al., 2016), all of which make a standard A+G method less than ideal. First, the two cues to be presented should ideally be matched in their reliability; participants should be about as precise with either cue. This is the situation in which the potential benefit of combination is greatest, and so the one in which the Bayesian optimal prediction is as different as possible from the alternative hypothesis that only one cue is being used. Second, it is also ideal to use a task for which the noise in perception/memory is approximately normally distributed around the target. This makes it possible to analyse precision (1/variance), which generally provides more statistical power than discrete nominal distributions. It also makes it possible to use simple, standard ways of predicting the optimal precision (Ernst & Banks, 2002). Third, it is ideal to use a situation where each participant can provide the highest possible number of trials, allowing for a within-subjects design. This makes it possible to calculate individual predictions for Bayesian optimal precision and compare these to individual measurements of precision with both cues. None of these three conditions are met in a standard A+G condition: the associative cue is more reliable than the geometric cue (Hermer-Vazquez et al., 2001; Lee, Winkler-Rhoades, & Spelke, 2012; Nardini et al., 2009), the errors are discretely distributed, and young participants will not generally tolerate much more than four trials in total.

Instead, we adapt a method from our previous studies (Negen, Heywood-Everett, Roome, & Nardini, 2018; Negen, Roome, Keenaghan, & Nardini, 2018). Virtual reality is used to make the trials faster and to make the task more engaging. Participants are shown a target being hidden among some landmarks. They then have their view blocked while their perspective changes. From this new perspective, the participant attempts to point where the target was hidden. On some trials, there is a pair of landmarks marking the North/South axis of the space. We refer to this as a NS trial. On other trials, there is a pair of landmarks marking the East/West axis. We refer to this as an EW trial. In the last kind of trial, both pairs of landmarks are available. We refer to this as a NSEW trial. This allows us to measure performance with two different cues (landmark pairs) in isolation and with both together. Participants included both adults and children (5-7 years), since Adaptive Combination is supposed to apply across the lifespan.

This design overcomes the usual problems described above. Since both cues are landmark pairs, they are matched in reliability. Responses on this kind of task are approximately normally distributed around the targets. Since more trials are tolerated, a within-subjects design is possible. This makes it a good way to test if reorientation cues are used together in a Bayesian manner.

## Hypotheses

Fourth (and finally), we detail the specific hypotheses and what predictions they make. For each of the three hypotheses, we first give a conceptual description in the top paragraph, followed by a bottom paragraph that lays out and justifies the specific empirical predictions about three outcome measures. Figure 1 is a reference guide for the different trial types and the empirical predictions of each hypothesis.

### Main Hypothesis: Adaptive Combination

As governed by Equation 1, the participant combines the information from the two landmark pairs in the optimal Bayesian fashion. This is a new extension (Xu et al., 2017) of the adaptive behaviour proposal (Cheng et al., 2013); it suggests that participants are not only taking account of which cues are available and which one is best, but also combining different cues while weighting them in line with Bayesian principles. This would be in line with how adults perform in many simple perceptual tasks (review, Pouget, Beck, Ma, & Latham, 2013). If this hypothesis fits children’s performance, that would break with the general pattern of children under 10 failing to show Bayesian reasoning (Burr & Gori, 2012) and warrant the exploration of a new theory of how Bayesian reasoning develops.

Figure 1 defines which trials are considered NSEW trials, NS or EW trials, and Near trials. The Adaptive Combination hypothesis predicts that precision in NSEW trials will be equal to the optimal Bayesian precision. In other words, the optimal Bayesian process should produce the optimal Bayesian precision. There is a simple and well-known formula used to predict the optimal Bayesian precision (Ernst & Banks, 2002). This hypothesis also predicts that NSEW accuracy will be better than NS or EW accuracy. This is predicted because the Bayesian process should always benefit from additional landmarks – NSEW trials have four landmarks, but NS or EW trials have only two. NSEW accuracy should also be better than Near accuracy for the same reason (Near trials also have only two landmarks).

### Alternative Hypothesis: Adaptive Selection

If participants do not use multiple cues with full Bayesian efficiency, they may still adopt a sensible strategy that constrains error while only using one landmark. Under Adaptive Selection, participants select the landmark nearest to the target, encode the target location against it, and ignore the other landmarks. In doing so, they improve average performance over just using a random landmark – but not as consistently as a Bayesian process would. This is more in line with older forms of the adaptive proposal (Cheng et al., 2013). It posits that children take account of which cue will be most useful and use this to guide which cue they use, but does not entail any combination of landmarks, i.e. Bayesian reasoning. This would be in line with previous results where children are capable of selecting the best single cues. For example, they tend to prefer visual cues for judging spatial relationships and auditory cues for judging temporal relationships (Gori et al., 2012). However, it would not allow for any new conclusions regarding Bayesian reasoning in development.

This hypothesis predicts that precision in NSEW trials will be worse than the optimal Bayesian prediction. In other words, a non-optimal non-Bayesian process should not lead to the optimal Bayesian precision. This hypothesis also predicts that accuracy in NSEW trials will be better than accuracy in NS or EW trials overall, since the NSEW trials will sometimes have a nearer (better) single landmark to select and use. For example, look at the top middle target in Figure 1. The nearest landmark, at the North, is visible on every NSEW trial. However, it is not present during half of the NS or EW trials. This should drive higher NSEW accuracy than NS or EW accuracy. However, Near accuracy should be equivalent to NSEW accuracy, since they both provide the nearest (best) possible landmark to select and use. For example, looking again at the top middle target, the North landmark is visible for all NSEW trials and all Near trials.

### Null Hypothesis: Random Cue Selection

On a trial with both landmark pairs, the participant chooses one landmark at random and encodes the target against it. The other landmarks are ignored. In essence, under this hypothesis, a NS or EW trial is a NSEW trial where we have done some of the random choosing for the participant. This would be similar to how children performed in a previous spatial task with two cues available, alternating in an unpredictable way between self-motion information and landmark information (Nardini et al., 2008). However, again, it would not allow for any new conclusions regarding early Bayesian reasoning.

This predicts that precision in NSEW trials will be worse than the optimal prediction. This is again because the non-optimal non-Bayesian process should not produce the optimal Bayesian precision. It also predicts that accuracy in NSEW trials will not be different from accuracy in NS or EW trials overall. For example, we can look at the top middle target in Figure 1 again. On a NSEW trial, we only expect them to use the North (best) landmark on one out of four trials. We would expect the same thing for NS or EW trials (two trials would have the North and South available, with the North selected on one trial). This hypothesis further predicts Near accuracy will be better than NSEW accuracy. For Near trials, we would expect them to use the North landmark two times out of four. In other words, in a Near trial, the lack of poor encoding choices should actually help participants if they are choosing encoding references randomly.

## Experiment 1

### Method

#### Participants

There were a total of 36 participants tested. Of these, 12 were adults (seven female). They ranged from 18 to 23 years old, with a mean of 20.9 years and a standard deviation of 1.25 years. The remaining 24 were children. Four did not complete the task, one because the headset was too large and three due to mood. Of the remaining 20 (four females), they ranged in age from five years and zero months to seven years and five months, with a mean of 6.1 years and a standard deviation of 0.6 years. All participants were recruited in the Durham, UK area. To the knowledge of the researchers, no children had been diagnosed with any perceptual or developmental disorder that might have affected task performance. The advertisements asked only for participants with normal vision or vision that could be corrected to normal with contact lenses. Adult participants (Psychology undergraduates) earned credits in a scheme where undergraduates participate in each other’s research projects. Children were given a small toy of their choosing. Written informed consent was obtained, either from the adults themselves or the parents of the children. Verbal assent was also obtained from the children themselves.

#### Apparatus

The study used WorldViz Vizard 5 software and the Oculus Rift headset. It also used an Xbox One controller. The virtual world (Figure 1) contained three major features situated around a 5m x 5m virtual space. First, there was a set of train tracks in a circle around the central space with a small cart. The cart could move around the tracks and had opaque shutters that could come up and down. The participant’s perspective was always from within the cart. Second, there was a set of four landmarks which could fade in and out of view. They were each unique and distinctive: black spheres, grey pyramids, red blocks, and blue cones. With the centre of the space at (0, 0), these were placed at the four cardinal points: (0, 3), (0, −3), (3, 0), and (−3, 0) in meters. Third, there were the diggers. These were the characters that played the game with participants. To make them more engaging to the children, they were given silly names and apparel. One digger, who had a moustache and wore a pipe hat, was named Digger T. Diggington III. The other digger, who wore a set of glasses with jewels and a large feather attached to a band around her head, was named Martha Diggington, Esquire. The 3D models for the Diggingtons had joints in the digging arm to their front so that they could be animated as digging a place for the target and then digging it back up. Fourth, there were the jewels. These served as targets to find. They were translucent blue (80% opacity) and fashioned after a round brilliant cut. There were no other landmarks or features in the environment that could be used to reorient (e.g. the skybox was uniform blue). The ground had a repeating sand texture at 20% opacity.

#### Procedure

The game began by allowing the participant to select the character they wanted to play with. The other character faded out of view. The first warmup trial began.

Each trial involved a series of four steps (Figure 2). First, the target was shown. The digger went to the target location. They stayed there for 3.5 seconds while an animation played of the target (jewel) being buried. The last 0.5 seconds involved the jewel going 3m into the air and moving straight down into the ground to make it as clear as possible exactly where it was.

**Figure 2.**
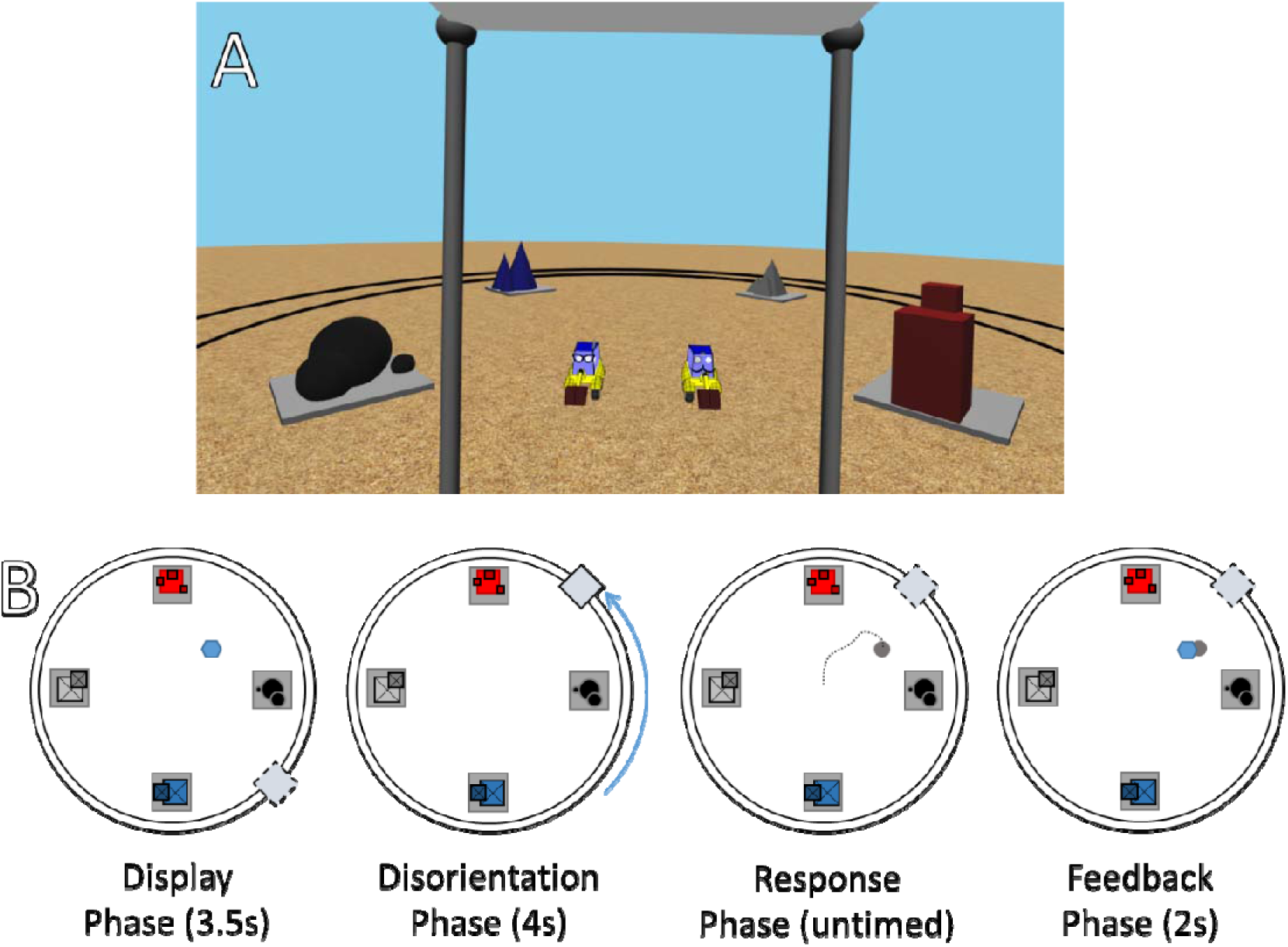
Methods for Experiment 1. (A) A first-person screenshot of the view within the experiment. Please be aware that the lenses of the Oculus Rift slightly distort the internal screen image, so the image given to it is distorted in the opposite way. For example, in the headset, it is clear that the red and blue landmark face each other directly; in the screenshot, they appear slightly offset. (B) First, the target (blue hexagon) was shown to the participant while they were in the cart (dashed box). Then the cart ‘closed’, blocking their view, and the participant was moved +90, −90, or 180 degrees around the track (black circles). Then the view was opened and the participant moved a grey cone to the point where they believed the target to be. Finally, feedback was given as to the correct placement. This could be done with either the North and South landmark (red and blue), the East and West landmark (grey and black), or all four.

Second, there was the disorientation. The opaque shutters on the cart moved up to block the participant’s view. Three seconds passed while a sound effect of a training moving played. The viewpoint changed. The shutters then lowered. This took a total of four seconds.

Third, there was the response. The participant used the joystick on the controller to move a large arrow with its tip on the ground within the 5m x 5m central space. There was a grey circle on the tip of the arrow with a radius of 75cm. When satisfied with the location, the participant pressed a button on the controller to enter their response. They were allowed as much time as they wanted, but younger participants were encouraged to take their best guess if they said that they did not know the right place.

Fourth, there was feedback. The digger moved over to the response location and played a 2s digging animation. If the response was within 75cm of the target, the jewel appeared out of the ground, a small ‘ding’ sound played, and the digger jumped up and down in a celebratory animation. If not, no jewel appeared, no sound played, and the digger turned towards the participant to play a ‘deflated’ animation. Over the course of 1s, their body widened along the ground plane by 20% while their height shrunk 20%. It then returned to normal shape over the next 1s. During this, a small blue circle flashed on the ground at the correct target location. When a button on the controller was pressed again, the next trial began.

The first five trials were considered warmup trials. These data are not analysed in any part of the results. During this time, the experimenters gave the children hints and explanations about the game. For remaining trials, participants were not given any extra information about the target location.

#### Stimuli and Trial Parameters

Target locations were on a 5×5 grid with 1m spacing. For example, there was a corner target at (2, 2), a center target at (0, 0), an off-center target at (0, −1), and a target in front of the West landmark at (−2, 0). For the five warmup trials, the targets were always (0, 0), (0, 2), (2, 0), (−2, 0), and (0, −2). After that, for adults, all 25 possible target locations were used. For children, to make the game shorter, only nine target locations were used: (0, 2), (−1, 1), (2, 1), (−1, 0), (0, 0), (1, 0), (−2, −1), (1, −1) and (0, −2). Each target was tested once with the East and West landmarks (EW trial), once with the North and South landmarks (NS trial), and once with all four (NSEW trial). The order of trials was random. This means, in total, that adults produced 75 analysed trials each and children produced 27.

The cart could travel either +90, −90, or 180 degrees around the track. This was done because the corners provided a good view of the target space where all four landmarks were visible but not obstructing the 5m x 5m response area. The amount of travel was chosen randomly on each trial. Each trial began wherever the last one ended.

#### Analysis Plan

To analyse these data, we plan to have four tests. First, just to confirm that the task was understood by participants, we will check that target locations and response locations were significantly correlated along both the x-axis and the y-axis. After this, responses will be excluded as outliers if they are more than 2.5 standard deviations in error away from the target.

To make the next three tests clear, we need to comment on accuracy, mean error, precision, and variable error. Some of the hypotheses are stated in terms of accuracy. To be more specific, we intend this as the mean error: the average distance between the target location and the response location, calculated along the 2D plane using the Pythagorean Theorem. Lower mean error indicates better accuracy. The other hypotheses are stated in terms of precision. To look at precision, we actually analyse variable error: the standard deviation of the response locations minus the target locations. As the variable error (standard deviation) of responses increases, precision decreases. Precision is conventionally defined as variable error raised to the power of negative two. Using variable error in the analyses, rather than precision, is standard practice in the cue combination literature (Ernst et al., 2016). This is because variable error tends to better approximate a normal distribution and tends to have a (much) less serious problem with sensitivity to outliers. In line with this, we will analyse and report variable error. Lower variable error indicates better precision. Conveniently, this means that a shorter bar denotes better performance in all of the bar graphs that will be shown. Since responses were along a 2D ground plane, there is a separate variable error along each axis of the space. We will use variable error to test specific predictions about reaching the optimal Bayesian precision; otherwise, we will use accuracy as a measure of performance.

For the second test, we will look at the Bayesian optimal variable error in NSEW trials, where all four landmarks were visible, versus observed variable error in NSEW trials. Adaptive Combination predicts that these will be equal. In other words, an optimal Bayesian process should produce the optimal Bayesian variable error. Adaptive Selection and Random Selection suggest that observed variable error with both cues will be worse than the optimal prediction. In other words, a non-optimal non-Bayesian process will fail to produce the optimal Bayesian variable error. For each participant, along each axis, for each trial type (NS, EW, and NSEW), we will calculate the variable error. For each participant, the optimal variable error is calculated with the equation (Ernst & Banks, 2002):

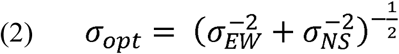

This comparison, as well as the next two, will be tested with paired t-tests. This second test conforms entirely with the standard method of testing for optimal Bayesian cue combination (Ernst et al., 2016).

Third, we need to test the accuracy in NSEW trials versus the accuracy in NS or EW trials, where only two landmarks were visible. Random Selection predicts that accuracy will be the same in NSEW trials versus NS or EW trials – under Random Selection, a NS or EW trial is just a NSEW trial where we have done some of the random selection for the participant. Adaptive Selection and Adaptive Combination predict that NSEW accuracy should be better than NS or EW accuracy, using the additional information in a NSEW trial to improve accuracy through either selecting the best single landmark (Selection) or via Bayesian cue combination (Combination).

Fourth, we need to test NSEW accuracy against Near accuracy. A trial is considered a Near trial if it is a NS trial or EW trial where a nearby single landmark is visible – at least as near as the nearest one in a NSEW trial with the same target (see Figure 1). This analysis proceeds on the assumption that accuracy at localising a target location using a landmark decreases as the target location gets further from the landmark (e.g. Negen, Roome, et al., 2018). Random Selection predicts that NSEW accuracy will be *worse* than Near accuracy, since the participant will sometimes randomly select one of the landmarks from the further (worse) pair to use on a NSEW trial. Adaptive Selection predicts that NSEW and Near accuracy will be equal, since participants complete a NSEW trial by only using the nearest landmark anyway. Adaptive Combination predicts that NSEW accuracy will be *better* than Near accuracy, since the Bayesian framework allows information from the further (worse) landmarks to be incorporated in a way that it still improves the responses.

### Results

Results strongly favour Adaptive Selection for both adults and children. For adults, the responses were correlated with the targets along the x-axis, r(898) = 0.83, p < .001, and the y-axis, r(898) = 0.80, p < .001 (Figure 3). Responses were excluded if they were more than 2.5 standard deviations away from the target (2.1m; 4.3% or 77 observations). Variable error was worse (higher) than the Bayesian optimal variable error along both the E/W axis, t(11) = −1.97, p = 0.038, and the N/S axis, t(11) = −2.76, p = 0.009 (Figure 4). Accuracy was better in NSEW trials versus the NS or EW trials, t(11) = t(11) = −3.02, p = 0.012 (Figure 5). Accuracy was not better in the NSEW trials versus the Near trials, t(11) = 0.21, p = 0.839.

**Figure 3.**
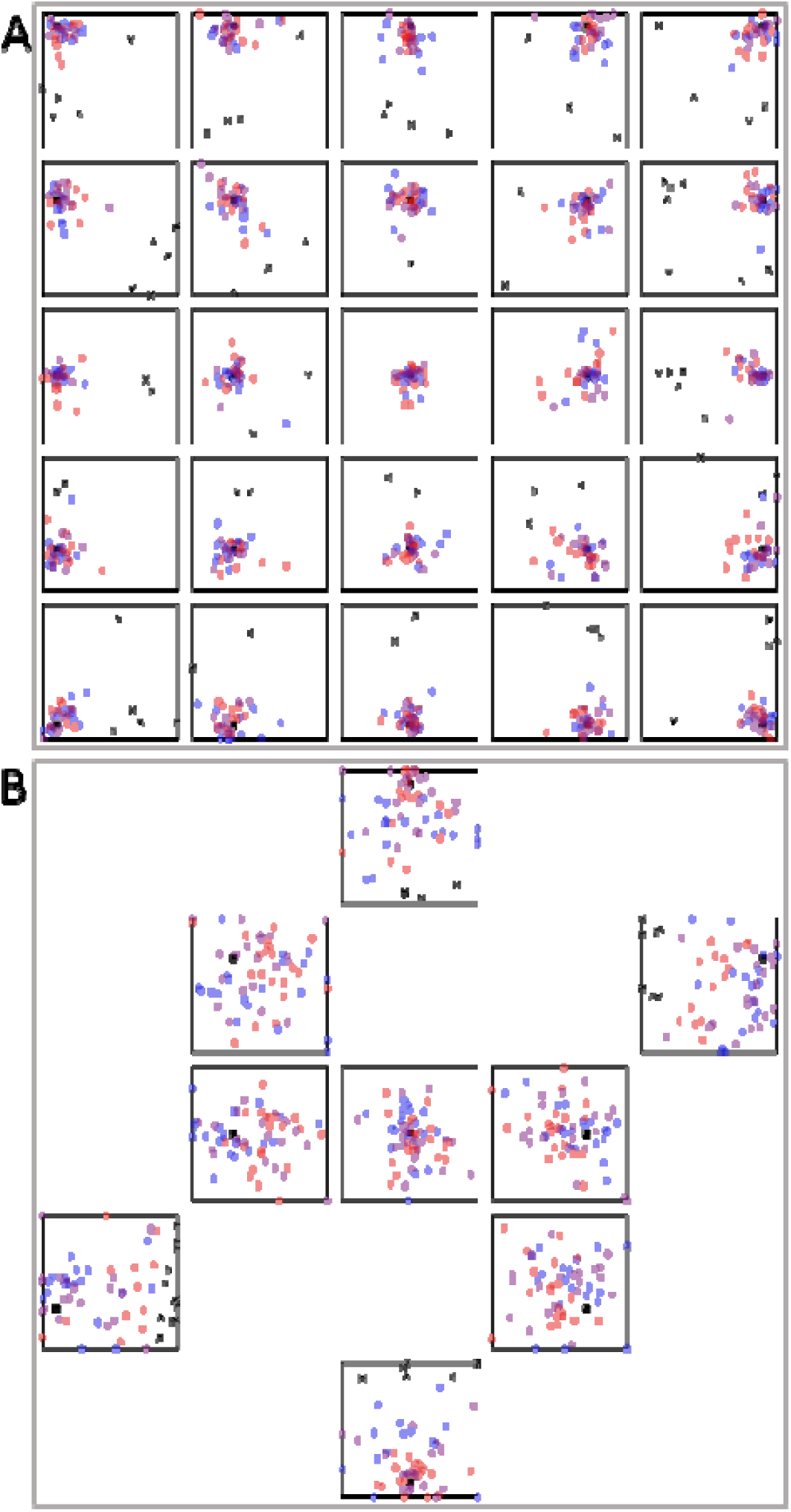
Adult (A) and Child (B) data from Experiment 1. Red dots are responses on NS trials, where the North and South landmark are visible. Blue dots are EW trials. Purple dots are NSEW trials. The black square is the target. Black crosses are excluded trials.

**Figure 4.**
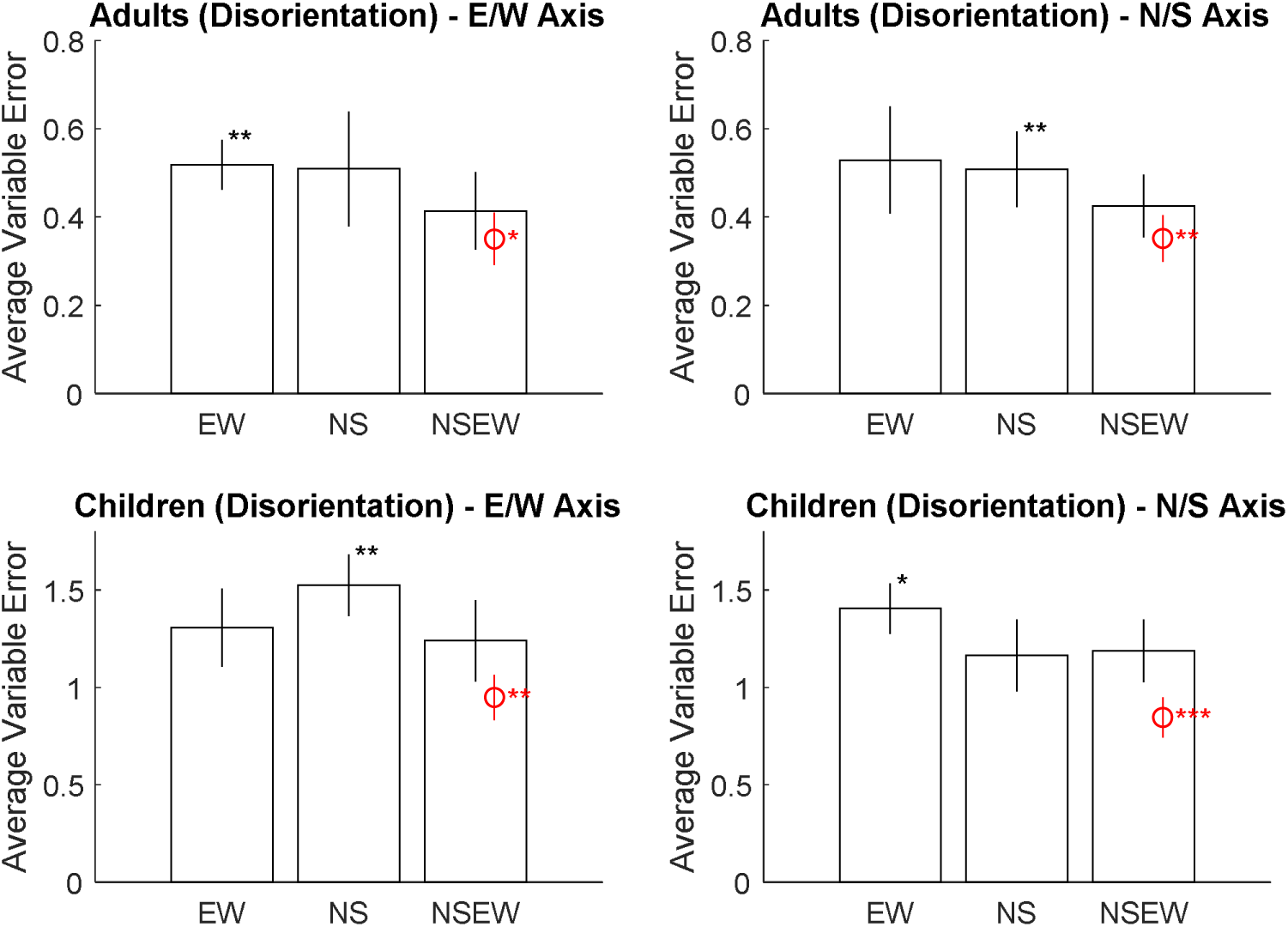
Average variable error broken down by trial type (x axis), participant group (top vs bottom panels), and axis of the space (left vs right panels). Error bars are 95% confidence intervals for the mean. Asterisks mark significant paired t-tests: *p<.05, **p<.01, ***p<.001. The red marking is the Bayesian optimal variable error. Both groups, along both axes, had significantly higher (worse) variable error than the Bayesian optimal variable error when shown all landmarks. This speaks against Adaptive Combination, but is consistent with either Adaptive Selection or Random Selection.

**Figure 5.**
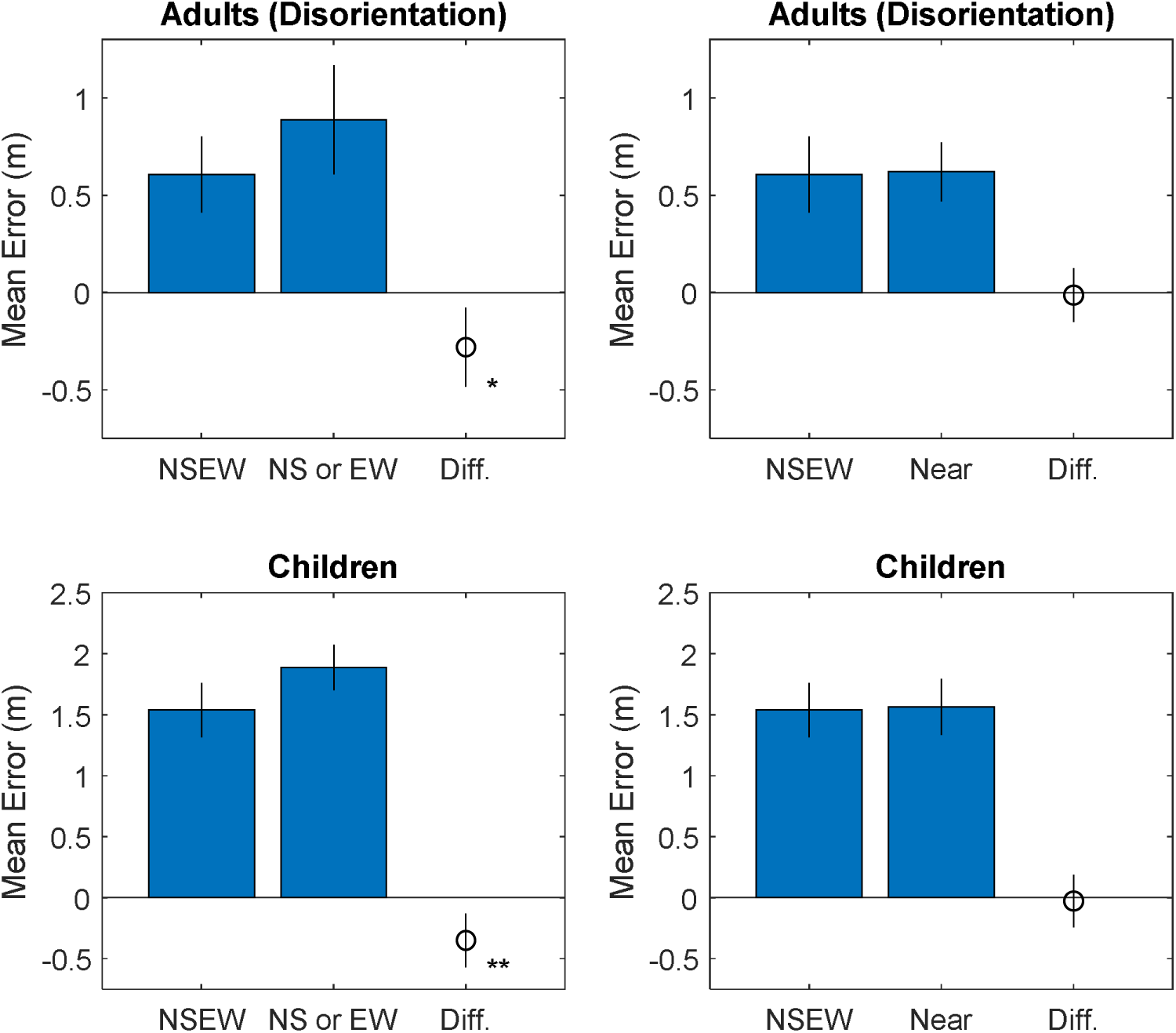
Accuracy compared with the NSEW trials, broken down by group (top versus bottom panels), trial type (x axis), and comparison trials (left versus right panels). Results favour Adaptive Selection, which predicts a difference versus NS or EW trials but not versus Near trials.

If Adaptive Combination were correct, we would not expect to see a difference between the optimal variable error and the observed variable error with both cues. We would also expect to see that NSEW accuracy was better than Near accuracy. If Random Selection were correct, we would not expect to see a difference between NSEW accuracy versus NS or EW accuracy. We would also expect to see that NSEW accuracy was worse than Near accuracy. Adaptive Selection correctly predicted that the variable error with both cues would be worse than optimal, that NSEW accuracy would be better than NS or EW accuracy, and that there would not be a difference between NSEW accuracy and Near accuracy.

For children, the pattern was the same (but with worse variable error and accuracy). The responses were correlated with the targets along the x-axis, r(537) = 0.28, p < .001, and the y-axis, r(537) = 0.31, p < .001 (Figure 3). Responses were excluded if they were more than 2.5 standard deviations away from the target (3.75m; 2.9% or 31 observations). Variable error was worse (higher) than the Bayesian optimal variable error along both the E/W axis, t(19) = −3.13, p = 0.003, and the N/S axis, t(19) = −3.87, p = 0.001 (Figure 4). Accuracy was better in NSEW trials versus the NS or EW trials t(19) = −3.30, p = 0.004 (Figure 5). Accuracy was not better in the NSEW trials versus the Near trials, t(19) = 0.25, p = 0.803. By the same logic as the adults, this favours Adaptive Selection.

### Interim Discussion

The results of Experiment 1 point towards Adaptive Selection for both adults and children. Adaptive Selection is a non-Bayesian process of selecting the best single cue and using it in isolation. For children under 10 years, this is in line with previous research regarding the use of multiple cues (Adams, 2016; Burr & Gori, 2012; Chambers et al., 2018; Dekker et al., 2015; Gori et al., 2012; Jovanovic & Drewing, 2014; Nardini et al., 2010, 2013, 2008; Petrini et al., 2014). Re-analysis of previous data agrees as well. This means that, in regards to the children, we now have a consistent and clear pattern of results. They likely do not use a Bayesian process in the classic geometric reorientation paradigm (see the re-analysis of A versus A+G conditions in *The Need for Additional Scrutiny*). They do not use a Bayesian process in the present paradigm. They do not use a Bayesian process when given landmark and self-motion cues (Nardini et al., 2008). Children under 10 generally do not use multiple cues in a Bayesian manner (Burr & Gori, 2012; though see Negen et al., 2019).

For adults, when considering both the present result and the previous literature, the overall pattern of results is somewhat disjointed and requires further examination. Adults can frequently use a Bayesian process in perception and memory (Pouget et al., 2013). It is not clear why adults would not have used a Bayesian process here. The next experiment is designed to see why this was occurring.

To isolate the variable preventing cue combination, we can closely compare Experiment 1 and a previous study that did find cue combination (Jetzschke, Ernst, Froehlich, & Boeddeker, 2017). Both studies used adults. Both studies used a virtual reality method. Both studies used multiple landmarks as the different cues. However, there are two differences. The previous study did not use an explicit disorientation procedure. Participants were led from a study location to a release location in a circuitous way, but with their eyes open and the landmarks always visible. This makes it difficult to trace the exact route back to the study location, but never particularly induces a sense of disorientation. This might be important because disorientation may induce specific neural processes that attend to specific spatial cues more than others (Cheng, 1986; Keinath et al., 2017; Knierim, Kudrimoti, & McNaughton, 1995). The previous study also used a homing task, asking participants to return to the homing location, rather than a recall task, asking participants to select where a target location was presented. This is potentially important because homing relative to landmarks can be completed in a completely egocentric fashion, just remembering a ‘snapshot’ of what the landmarks looked like from the studied home viewpoint (Stürzl, Cheung, Cheng, & Zeil, 2008). The task here requires a completely allocentric strategy. Experiment 2 is therefore as similar as possible to Experiment 1, except it also removes the disorientation aspect; it disrupts egocentric vectors to the targets in a way that does not disorient the participant. If cue combination is observed, then the disorientation is likely preventing cue combination. If not, then the difference is likely due to the task itself (homing versus recall) and its implications in terms of egocentric vs allocentric reasoning.

## Experiment 2

Experiment 2 is an experiment done solely with adults, as similar as possible to Experiment 1, but without disorientation. This was done to test the hypothesis that adults will combine cues in allocentric spatial tasks without disorientation, but not allocentric spatial tasks with disorientation.

### Method

The method was as similar as possible to Experiment 1, except without disorientation (Figure 6). In short, we spun the target and landmarks instead of the participant. To make this possible, the virtual environment was altered. The target area and the landmarks were raised onto a circular pedestal. The pedestal had identical markers placed around its edge. The ground near the pedestal also had identical markers. There was also a grey half-sphere that could appear over the top of the pedestal, blocking all vision of the target area and the landmarks. The participant’s viewpoint was set back another 2m so that they could see the spinning platform and the stationary ground around it, making it clear that the platform specifically was spinning (and not the participant moving around it). After being shown the target, the participant was not moved or turned in any way. Instead, the grey half-sphere covered the pedestal. The pedestal spun rapidly and erratically for three seconds, making it impossible to track the target egocentrically. The grey half-sphere was removed. The participant then attempted to point to the target location. This requires the participants to use the landmarks, which is the same as Experiment 1. One might think of this as a local or intrinsic allocentric frame. However, it induces no sense of disorientation. Beyond this, the experiment was the same as Experiment 1.

**Figure 6:**
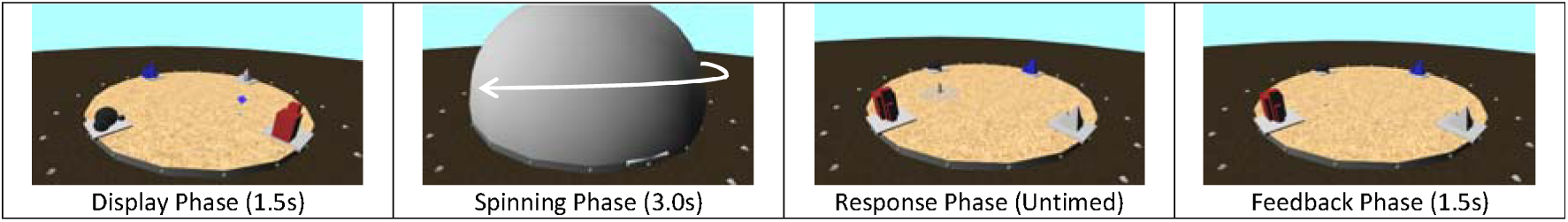
Methods for Experiment 2. Participants were shown the target. The target and landmarks were covered and then spun rapidly and erratically. The cover was removed and the participant would then indicate the target from memory.

### Results

Results again favour Adaptive Selection. The responses were correlated with the targets along the x-axis, r(898) = 0.91, p < .001, and the y-axis, r(898) = 0.90, p < .001 (Figure 7). Responses were excluded if they were more than 2.5 standard deviations away from the target (1.5m; 2.6% or 46 observations). Variable error was worse than the Bayesian optimal variable error along both the E/W axis, t(11) = −2.15, p = 0.028, and the N/S axis, t(11) = −1.93, p = 0.040 (Figure 8). Accuracy was better in NSEW trials versus the NS or EW trials, t(11) = −7.15, p < .001 (Figure 9). Accuracy was not better in the NSEW trials versus the Near trials, t(11) = 0.30, p = 0.772. All of these patterns are the same as Experiment 1.

**Figure 7.**
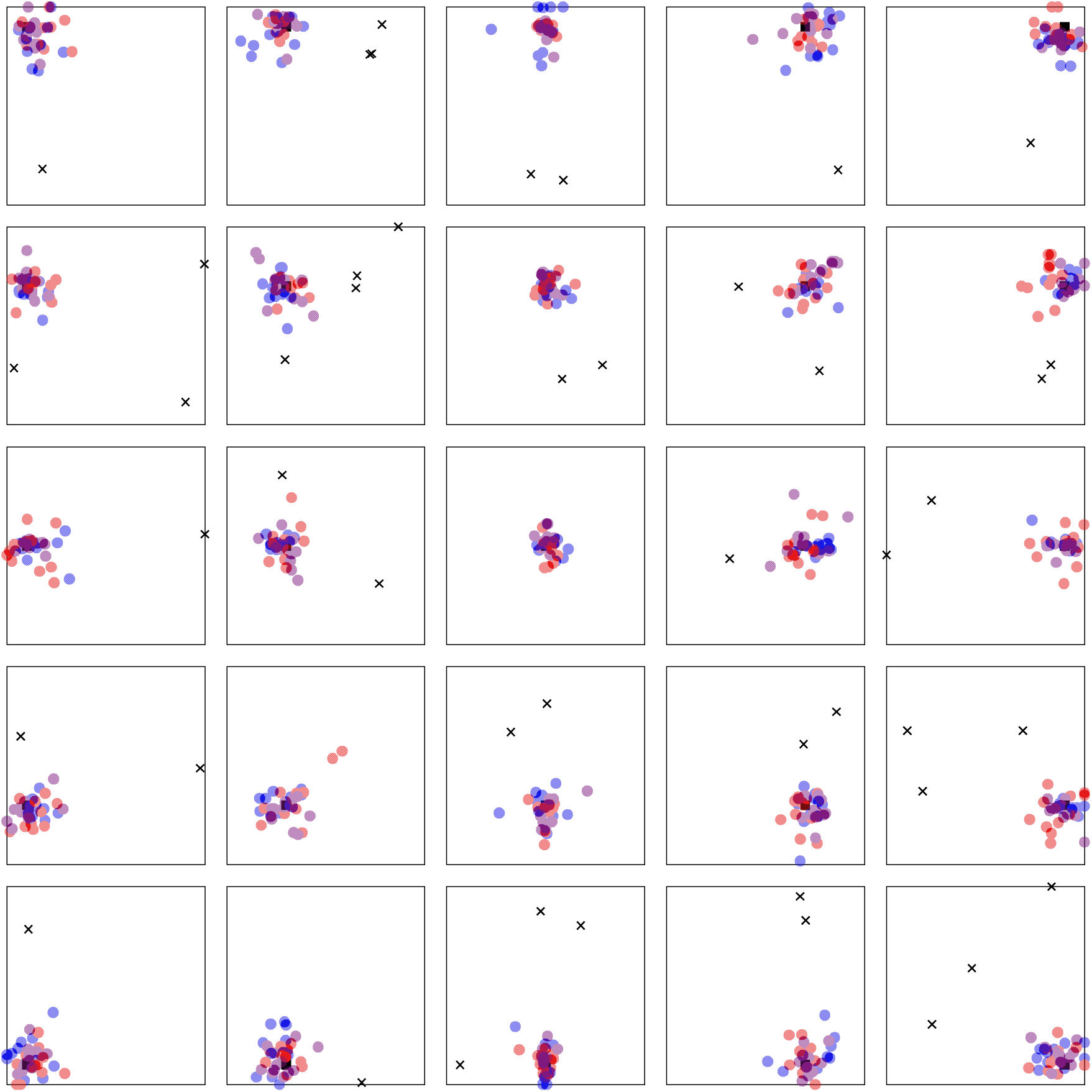
Adult data from Experiment 2 (without disorientation). Red dots are responses on NS trials, where the North and South landmark are visible. Blue dots are EW trials. Purple dots are NSEW trials. The black square is the target. Black crosses are excluded responses.

**Figure 8.**
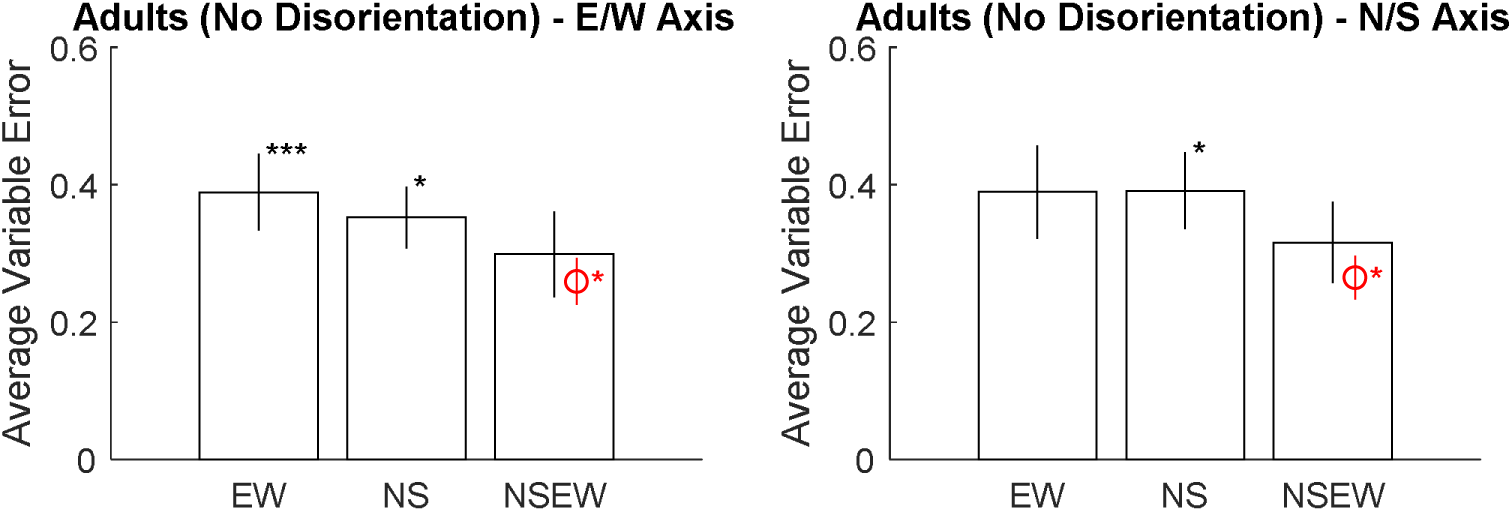
Average variable error broken down by trial type (x axis) and axis of the space (left vs right panels). Error bars are 95% confidence intervals for the mean. Asterisks mark significant paired t-tests: *p<.05, **p<.01, ***p<.001. The red marking is the optimal prediction. Along both axes, participants had significantly higher variable error than the optimal prediction when shown all landmarks. This speaks against Adaptive Combination, but is consistent with either Adaptive Selection or Random Selection.

**Figure 9.**
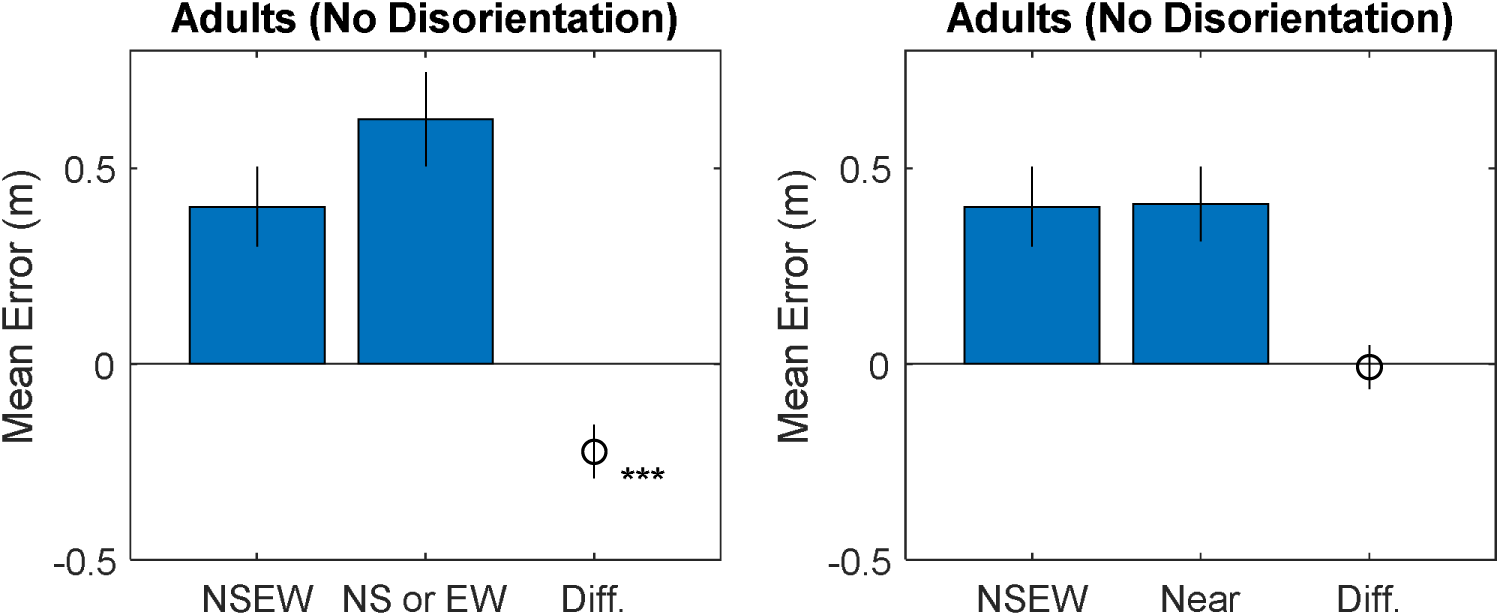
Accuracy compared with the NSEW trials, broken down by trial type (x axis) and comparison trials (left versus right panels). Results favour Adaptive Selection, which predicts a difference versus NS or EW trials but not versus Near trials.

While the results are the same as the adults in Experiment 1 in terms of favouring Adaptive Selection, the lack of disorientation did lead to better overall performance. In a 2 (Disorientation vs No Disorientation) by 2 (NSEW vs NS or EW) ANOVA, using mean error as the dependent variable, there was a significant effect of disorientation, F(1,44) = 7.47, p = .009, η^2^ = .124. Similarly, in a 2 (Disorientation vs No Disorientation) by 2 (NS axis vs EW axis) by 3 (NS, EW, or NSEW landmarks visible) ANOVA, using variable error as the dependent variable, there was a main effect of disorientation, F(1, 139) = 31.29, p < .001, η = .170, with worse (higher) variable error after disorientation.

### Interim Discussion

Experiment 2 was done to see if the difference in results between adults in Experiment 1 and a previous study (Jetzschke et al., 2017) was due to the use of disorientation in Experiment 1. Since results were like Experiment 1 (i.e. not showing cue combination), but Experiment 2 did not involve disorientation, this hypothesis seems unlikely. Instead, this isolates a more fundamental aspect of the tasks: here, participants had to use landmarks in a local allocentric frame to recall locations, whereas the previous study asked participants to return ‘home’ in a way that allows egocentric snapshots to be useful. Other than this, Experiment 2 and the previous study both tested adults, used virtual methods, did not disorient participants, and used multiple landmarks as the cues.

## General Discussion

Both experiments point strongly towards Adaptive Selection, a non-Bayesian process of selecting the most useful landmark and using it isolation. They point away from Adaptive Combination, a Bayesian process. Specifically, the Bayesian predictions about precision were consistently violated. They also point away from Random Selection, a non-Bayesian process of selecting a landmark to use at random. Specifically, accuracy was better than we would expect from using a random landmark. In contrast, results are consistent with all three predictions if participants are just encoding the target location against the nearest single landmark. We interpret this to mean that landmarks are not used together in a Bayesian fashion to recall locations, at least in a situation where egocentric relations have been disrupted; instead, people use the nearest available landmark to code locations. This provides an immediate theoretical point: that the Adaptive Combination model, taken as a general theory of how multiple cues are used to reorient, is not as broadly applicable as one might have hoped. We propose considering the older Adaptive Selection model, which still allows young children to use superior cues *in place of* inferior cues when both are available, but not to use superior cues in Bayesian combination with inferior cues.

To aid in interpretation, we need to point out a few things about the current study. Our focus was not particularly on the way that boundaries, including rectangular boundaries, are used to reorient. Instead, the goal of the design here was to find a situation where the predictions of a Bayesian cue combination model for reorientation could be clearly confirmed or discredited. Under our reading, Adaptive Combination is intended to be a flexible framework for the way that any set of valid cues are used to reorient – not just rectangular enclosures. To make the predictions of this framework as clear as possible in the present study, we used pairs of landmarks as cues. The results here speak against the general form of the Adaptive Combination model as a way for any reorientation cues to combine for allocentric recall. For a researcher who is specifically interested in the use of rectangular enclosures, rather than a general theory of how reorientation happens, the new data presented here have a more modest interpretation. It could still be the case that other cues are used in a Bayesian fashion to reorient, perhaps even at young ages. We suggest holding off on that conclusion unless and until more evidence for it is found.

We should also point out that a variation without disorientation did not appear to alter results. In other words, these results do not appear strictly limited to reorientation. Instead, they appear to apply to situations where egocentric relations are broken. It does appear that using landmarks to reorient is a non-Bayesian process, but it may make more sense to describe this in terms that are more general: landmarks are not used in a Bayesian process to recall locations when the use of the allocentric frame is forced.

Our re-analysis of previous data also suggests that geometric and associative cues are not combined in a Bayesian fashion by young children, but here we have to be more tentative. In the Introduction, we re-analysed previous data to compare performance in an associative-only reorientation task versus an A+G (one uniquely colored wall in a rectangle) reorientation task. No difference was found. This does not fit well with the idea that the associative cue’s information is being combined in the optimal, Bayesian manner with the geometric cue’s information. Instead, it suggests that the associative cue’s information is used in isolation. However, this analysis is far from ideal. For example, it uses between-subject data. In our view, this specific question remains open.

It should also be pointed out that Adaptive Combination and Adaptive Selection can make nearly identical predictions in the right circumstances. For example, suppose a child is given a very strong associative cue (e.g. a very salient and non-symmetric picture on one wall) and a very weak geometric cue (e.g. a rectangular boundary with a length of 2m and a width of 2.05m). Adaptive Selection would select the associative cue and the child would perform as if they only had the associative cue. Adaptive Combination would weight the two cues together according to Equation 1 – but since the geometric cue is much weaker, it would receive negligible weight, and the results would not be measurably different to those based on using the associative cue alone. In general, the two theories make very similar predictions in any situation with one dominant reorientation cue. Differences can only become clear when there are multiple reorientation cues with comparable reliability.

As far as we are aware, the present interpretation of an A+G condition is novel. In the developmental literature, it is well established that young children can use purely geometric cues to reorient (Lee, 2017). In interpreting the results of an A+G condition, the usual question has been whether the associative cue is used in concert with the geometric cue (Cheng et al., 2013; Hermer & Spelke, 1994). It could be the case that the associative cue is used in isolation while ignoring the geometric cue – at least in situations with a relatively large room. (In a small room, in contrast, it is well established that performance is similar to only having the geometric cue.) It may be possible to test an exclusive reliance on the associative cue more directly in the future, but it would require some significant methodological innovations. Ideally, the same participants would complete a large number of A, G, and A+G condition trials. It is not obvious how to make the standard paradigm into something that will be tolerated by young children for significantly longer. Further, details of the method would need to be adjusted somehow to make A performance better-matched to G performance – perhaps by reducing the contrast of the associative cue and exaggerating the ratio of the rectangle’s lengths. Further, and perhaps most difficult, it is not clear how this kind of paradigm would differentiate Bayesian reasoning from other simpler models. For example, the information from the geometric and associative cue could be combined through conjunctive logic (e.g. search until finding a target that agrees with both remembered cues) rather than probabilistic Bayesian reasoning. This would also predict that A+G performance would be better than A performance. It may ultimately be more fruitful to move to new paradigms.

We should emphasize that Adaptive Combination is a recent extension of the adaptive behaviour position; rejecting Adaptive Combination does entail rejecting all adaptive explanations of how young children reorient. It should not be interpreted to mean that a modular theory (Hermer & Spelke, 1994), the usual contrast to an adaptive theory, should be preferred. It would require a very different kind of experiment to potentially show evidence for non-adaptive and modular cognition (Lee, 2017; Lee & Spelke, 2008, 2010). Instead, Adaptive Selection is more in line with the versions of adaptive theories proposed before Adaptive Combination (Cheng et al., 2013). In addition, while our results speak against Adaptive Combination as a general theory of spatial reorientation, it remains possible that it does apply to some situations – for example, recall that can include an egocentric process of homing (Jetzschke et al., 2017; Stürzl et al., 2008), or recall using other kinds of spatial cues (although see above on difficulties of testing these in a Bayesian framework).

One interesting future direction would be to test the efficiency of cue selection in a setting where there are not just landmarks. If one landmark is further than another is, it is fairly clear that it will be less useful for encoding. If a young participant is asked to choose among a more diverse set of cues (e.g. including a linguistic cue), it is not yet known if they will consistently select the most useful cue.

### Towards a More General Theory

Here we outline how these results may drive us towards a more general theory of how reorientation happens with multiple cues. The present study provides an immediate empirical conclusion: children and adults select the nearest landmark to use in isolation for encoding targets during an allocentric reorientation task. A larger interpretive framework will require more research. To move this forward, we will sketch one (of many) plausible models to pursue and test further. In short, we consider that egocentric spatial information may typically be treated in a Bayesian manner after a certain point in development, sometime in middle childhood; allocentric information, across the lifespan, may instead be processed with more idiosyncratic non-Bayesian heuristics.

That idea has three parts. First, egocentric spatial information is typically used in a Bayesian manner by adults. This fits in a variety of very simple perceptual tasks where participants are asked to make judgements about locations. For example, adults can combine a spatialized sound and a noisy visual cue to judge horizontal location in an egocentric frame (Battaglia, Jacobs, & Aslin, 2003; Gori et al., 2012). This also applies to newly-learned skills that signal egocentric distance, such as an echolocation-like skill taught over the course of a few hours (Negen, Wen, Thaler, & Nardini, 2018). This further applies in a navigation task where the two cues are vestibular and proprioceptive (Frissen, Campos, Souman, & Ernst, 2011). Adults also rapidly learn egocentric (sensorimotor) prior distributions and use them in a Bayesian fashion as well (Bejjanki, Knill, & Aslin, 2016; Berniker, Voss, & Kording, 2010; Chambers et al., 2018; Körding & Wolpert, 2004; Kwon & Knill, 2013; Narain, van Beers, Smeets, & Brenner, 2013; Sato & Kording, 2014; Tassinari, Hudson, & Landy, 2006). In practice, this means that they learn where targets tend to be and bias their responses towards the places they tend to be most often. Finally, adults tend to adjust their search strategy in a visual search task when there is an uneven distribution of targets in an egocentric sense (Jiang & Swallow, 2013, 2014; Smith, Hood, & Gilchrist, 2010). This can be viewed as using egocentric prior distributions to affect decision-making.

Importantly, one could view self-motion and landmark information in a homing task as egocentric information. These are the cues and the task used in a series of studies where adults were fit well by a Bayesian model (Bates & Wolbers, 2014; Chen, McNamara, Kelly, & Wolbers, 2017; Nardini et al., 2008; Sjolund, Kelly, & McNamara, 2018; Zhao & Warren, 2015). The self-motion information could be viewed as an egocentric vector to the goal that is updated by perception of own movement. The landmark information, in this case, could be like an egocentric ‘snapshot’ of how the landmarks looked at the target (home) location (Cheung, Stürzl, Zeil, & Cheng, 2008; Stürzl et al., 2008). In other words, while landmarks are usually thought of as allocentric information, the specific way that landmarks looked from a previous home location could be stored in an egocentric format. This makes this finding fit with the idea that adults use egocentric information in a Bayesian fashion.

Of course, recent research has shown that this also faces some limits and suboptimalities (Rahnev & Denison, 2018) – many Bayesian processes are distorted under certain circumstances. For example, in one study, adults integrated multiple repeats of the same audio localization signal with lower-than-Bayesian efficiency (Jones, 2018). It is not clear exactly why this occurred, but adults’ performance was much nearer to optimal Bayesian integration when the signals were not exact repeats of each other. We should emphasize that it may be typical for egocentric information to be processed in a Bayesian way, but it will not be universal.

Second, egocentric spatial information is not used in a Bayesian manner by young children. This would explain their difficulty in making egocentric spatial judgements with an audio and a visual cue (Gori et al., 2012), difficulty combining self-motion and landmark information during a homing task (Nardini et al., 2008), and their difficulty in learning prior spatial distributions (Chambers et al., 2018). This fits more generally with the pattern of difficulty with Bayesian reasoning in a wide variety of settings (Burr & Gori, 2012).

Third, allocentric spatial information is not used in a Bayesian manner. This fits with all the findings here. Instead, what the participants did here appears to involve focusing on ‘just enough’ of the allocentric spatial relations to uniquely encode the target location in principle. It could be that once a task involves allocentric computations, capacity for the number of cues that can be attended to becomes a major bottleneck. At that point, it may be more advantageous to focus attention on the way that a target relates to a single nearby landmark than to spread attention across an entire scene. This also fits with visual search patterns in an allocentric frame. Participants do not tend to use the prior distribution to adjust their search strategies (Jiang & Swallow, 2013, 2014; Smith et al., 2010). This is, again, ‘just enough’ – learning the prior distribution does not affect the participant’s ability to complete the task; it only makes it faster to do so. It may be that allocentric reasoning is too slow for it to affect the way the visual search task is completed. In general, the complexities of the representations required for allocentric reasoning may be too slow and too costly to be a good application of Bayesian reasoning. That may be reserved instead for mature egocentric reasoning.

### Conclusion

The present study was designed further test predictions from the adaptive cue combination model of human spatial reorientation, understood here as a general model of how multiple cues are used to retrieve vectors to goal locations after losing one’s sense of heading and placement in a space. The results suggest that this theory needs modification since the optimal Bayesian predictions were consistently violated. Instead, response patterns were more consistent with a heuristic of only using the nearest single landmark (ignoring other landmarks rather than combining their information in a Bayesian fashion). Further, results are similar if egocentric relations are disrupted through a method without disorientation. As a sketch of a broader theory for further testing, we suggest that egocentric information may typically be used with Bayesian efficiency after middle childhood (emerging from roughly 7 to 12 years depending on task details), but that allocentric information is processed using non-Bayesian heuristics even into adulthood.

## Supporting information

Raw Data

## Acknowledgements

This work was supported by grant ES/N01846X/1 from the Economic and Social Research Council of the United Kingdom. This project has received funding from the European Research Council (ERC) under the European Union’s Horizon 2020 research and innovation programme (grant agreement No. 820185).

